# Response to “Expression reduction of biallelically transcribed X-linked genes during the female human preimplantation development”

**DOI:** 10.1101/701227

**Authors:** JC Moreira de Mello, MD Vibranovski, LV Pereira

## Abstract

Reinius and Sandberg (2019) have recently shown that X-linked balletic gene expression decreases during human preimplantation embryonic development, corroborating their previous hypothesis of X dampening as a mechanism of dosage compensation. Here we demonstrate that their analyses are based on biologically false premises, thus not demonstrating the phenomena of X dampening in humans.

## INTRODUCTION

In our previous article^1^ we argued that in order to test the hypothesis of dosage compensation by X chromosome dampening during human preimplantation development, proposed by Petropoulos et al. (2016)^2^, one should analyze expression of only X-linked genes with biallelic expression at each stage. Now Reinius and Sandberg (2019)^3^ present their analysis of expression of biallelic X-linked genes throughout development using our criteria of calling genes mono or biallelic.

## RESULTS AND DISCUSSION

However, the new data the authors present (Figures 2–4) are based on a biologically erroneous presumption: that if an X-linked gene is biallelically expressed in one cell of an embryo, this gene can be considered as biallelic in all cells of all embryos at that same stage. We had demonstrated that this is not the case with our previous analysis of allele-specific gene expression in human embryos, where we detect the same gene as monoallelic in some and biallelic in other cells of the same embryo (Figure 2, Moreira de Mello et al., 2017^1^; Figure 1 below).

**Figure 1:**
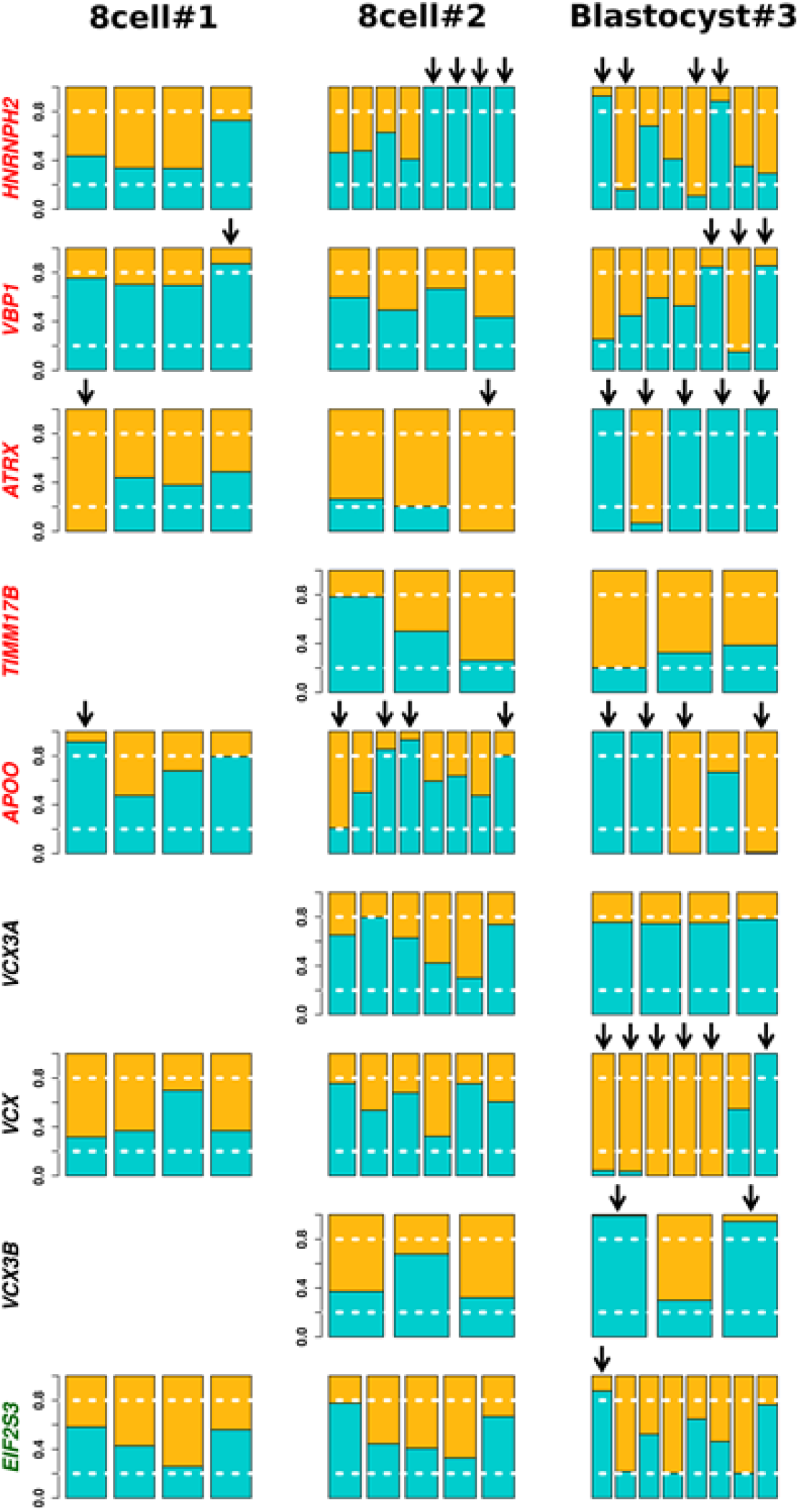
Allele-specific expression of X-linked genes at 8-cell and blastocyst stages. Cells are grouped by embryo where each bar represents a single cell. Alellic expression pattern was detected by informative SNPs in each embryo. Allelic relative expression ratios ≤ 0.2 or ≥ 0.8 were considered as monoallelic expression (white dotted line). Relative expression of reference and alternative alleles in blue and orange, respectively. Arrows point to cells presenting monoallelic expression. Y axis: relative expression ratio. Genes in red and green were described as subjected to and escaping XCI, respectively; those in black have no information^4^.

**Figure 2:**
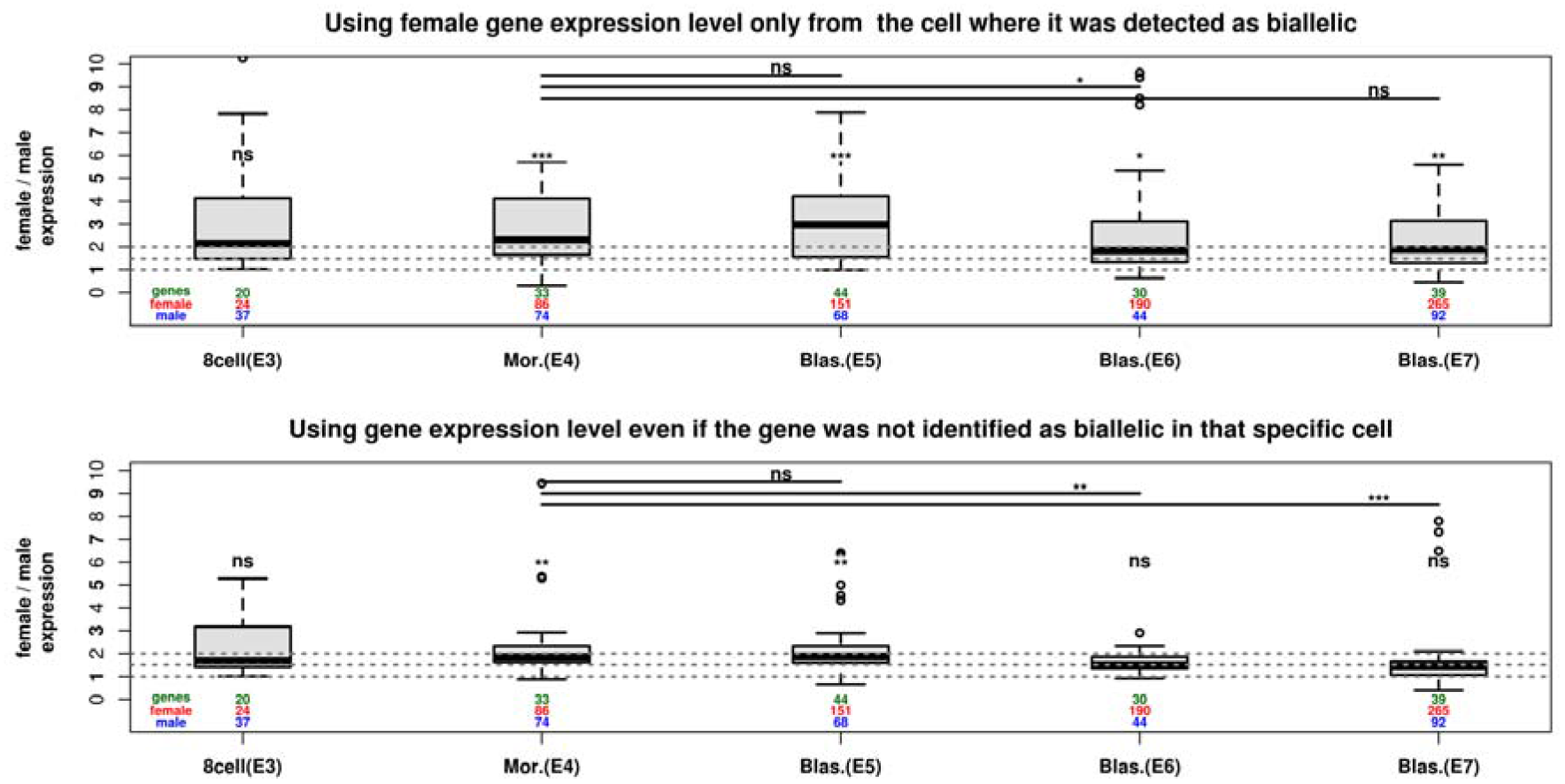
Distributions of female-to-male ratio of expression of X-linked genes biallelically expressed in females. Top panel: gene expression taken only from cells where biallelic expression was detected. Expression levels differ between females and males at morula and blastocysts stages. Despite the borderline difference between E4 and E6 (P-value = 0.045), there are no differences on female-to-male expression ratio among developmental phases, indicating no down-regulation of female X-linked biallelically expressed genes. Bottom panel: gene expression from all cells if a gene showed biallelic expression in any cell of that stage. Expression levels do not differ between females and males at E6 and E7 blastocysts stages, and female-to-male ratio of expression changes significantly from morula to blastocysts. Differences between medians of males and females in each stage are indicated above the respective boxplots. Differences in female-to-male ratios between stages are shown. Numbers of genes (green), of female (red) and of male (blue) cells analyzed are indicated for each stage. Wilcoxon tests: P-value (*) < 0.05; (**) ≤ 0.01; (***) ≤ 0.001; (ns) not significant.

Reinius and Sandberg (2019)^3^ incorrectly claim to have reproduced our analysis and reached a different result. In all our analysis of biallelically expressed genes we assumed nothing and used data only from those genes really detected as biallelic in each cell. In Supplementary Table 1 we present the list of biallelic genes per cell in one embryo of each developmental stage using Petropoulos et al^2^ dataset. Our analysis indicates lack of dosage compensation of biallellic genes by E7 (Figure 3D in Moreira de Mello et al^1^ reproduced here as Figure 2, upper panel), thus arguing against X dampening. If we make the calculations as they did, assuming a gene detected as biallelic in one cell is biallelic in all cells from all embryos in the same stage, indeed we detected dosage compensation of “biallelic genes” by E7 (Figure 2, lower panel). However, that latter strategy is biologically flawed as explained above.

The authors go on to make another puzzling assumption: that if a gene is biallelically expressed at E7, it can be assumed to be biallelic at all cells of all embryos from earlier stages. That would be a reasonable assumption if one were following gene expression of the same embryo throughout development. However, this is not the case – at each stage different sets of embryos are being analyzed. Again, Figure 1 (below) shows how heterogeneous allele-specific expression of a given gene can be within an embryo and among embryos at different stages.

Interestingly, their own data from 2016 had shown this heterogeneity within embryos. In Figure 6 of Petropulos et al.^2^, one can see how allele-specific gene expression varies within embryos, with genes being monoallelic in some and bialllelic in other cells of the same embryo. Interestingly, some genes, including *XIST* (*e.g.* embryos E6.6, E6.9), are monoallelic in more that 50% of the cells in some of the embryos, although the authors did not mention that in the text. Moreover, some genes are mostly biallelic in E7 embryos but monoallelic in many cells of embryos at earlier stages (*e.g.* PRPS: E7.16 vs E6.9; *MPP1*: E7.16 vs. E5.16). Therefore, results from Figures 3B and 4C-D (Reinius and Sandberg, 2019) are based in a biologically false premise.

In conclusion, we understand the potential limitations of analyzing allele-specific expression from scRNAseq data, especially when the sequencing depth is low (Moreira de Mello *et al*^1^, Supplementary Figure S2). For that reason, in our work we used complementary strategies in two different sets of scRNAseq data to address the issue of X-linked dosage compensation in human preimplantation embryos. And although our results indicate an early and heterogeneous process of XCI at the blastocyst stage and argue against dampening, the one thing we agree with Reinius and Sandberg (2019) is that further work must be done with better scRNAseq data from human embryos.

---

**Supplemental Table 1**:list of X-linked genes biallelically expressed (biallelic) in each cell of embryos E3.45, E4.9, E5.43, E6.6 and E7.12 (2). -: not determined.

## References

1. Moreira de Mello, J.C.M., Fernandes, G.R., Vibranovski, M.D. & Pereira, L.V. Early X chromosome inactivation during human preimplantation development revealed by single-cell RNA-sequencing. Scientific Reports 7(2017).

2. Petropoulos, S. et al. Single-Cell RNA-Seq Reveals Lineage and X Chromosome Dynamics in Human Preimplantation Embryos.Cell.165, (2016).

3. Reinius, B. and Sandberg, R. Expression reduction of biallelically transcribed X-linked genes during the female human preimplantation development. BioRxiv (2019).

4. Balaton, B. P., Cotton, A. M. & Brown, C. J. Derivation of consensus inactivation status for X-linked genes from genome-wide studies. Biol Sex Differ 6, (2015).

